# Cryo-electron microscopy ensemble optimization using individual particles and physical constraints

**DOI:** 10.64898/2025.12.02.691891

**Authors:** David Silva-Sánchez, Erik H. Thiede, Roy R. Lederman, Pilar Cossio

## Abstract

Biomolecules are inherently dynamic, and understanding their conformational ensemble distributions is essential for understanding their dynamics and biological roles. Cryo-electron microscopy (cryo-EM), a technique that images individual biomolecules frozen in a thin layer of amorphous ice, has emerged as a leading method for determining the structure of biomolecules at atomic resolution. Recent advances in cryo-EM reconstruction have made significant progress in determining structure in heterogeneous conformational landscapes. In contrast to reconstruction, a different class of techniques has been used to infer population weights, referred to as ensemble reweighting. These methods have yet to be generalized to infer structural heterogeneity simultaneously. Here, we present a method for *cryo-EM ensemble optimization* that directly infers the optimal set of structures and their associated population weights from cryo-EM images using Bayesian optimization techniques. Our method iterates between optimizing the structures and weights using a likelihood defined in terms of cryo-EM particle images (not reconstructions) and projecting onto the domain of a physical prior through an approach inspired by projected gradient descent. We test the method on several systems, ranging from a four-atom toy model to a large protein system with real cryo-EM data. We find that our approach successfully recovers the structures and their associated weights across a wide range of experimental conditions, even when the number of structures does not match the actual number of metastable states. Our method paves the way for cryo-EM ensemble optimization of flexible biomolecules exhibiting complex, multimodal conformational landscapes.

## 1 Introduction

To perform their biological functions, biomolecules shift between multiple conformational states. Some of these states are stable and highly populated, while others are short-lived but still critical for function. The probability distribution of each state, known as the ensemble distribution, reveals key thermodynamic properties, including activation barriers and rare intermediate or transition states that shed light on the molecular mechanism. Understanding this distribution is crucial for studying biomolecules, but measuring it—especially at high resolution across the whole conformational space—is exceptionally challenging.

Over the last decade, cryogenic electron microscopy (cryo-EM) has emerged as a powerful technique for studying the structure of biomolecules at atomic resolution and their conformational changes [1]. In cryo-EM, samples are flash-frozen and imaged using an electron microscope, preserving their native state. This process yields millions of noisy 2D projections of individual molecules embedded in the vitrified sample (micrograph), from which individual particle images are extracted for analysis. Each particle image captures a molecule in a different orientation and potentially a different conformation. Because flash freezing preserves the range of states present in solution, cryo-EM particles naturally sample conformations from the ensemble distribution [2, 3]. However, to achieve high-resolution reconstructions, particles are subjected to extensive 3D classification under the assumption that all particles within a given class adopt identical conformations [4]. Conformational-variability methods generally rely on a consensus reconstruction derived after 3D classification, and seek to model continuous motions around the consensus map [5–16].

Incorporating physical constraints can substantially improve the recovery of physically realistic structures from noisy cryo-EM maps. Integrative structural modeling combines physics-based simulations with experimental data by adding a data-driven energy term to the physical energy (force field), penalizing deviations from the measured observations. Molecular dynamics flexible fitting [17] was the first method to address this in cryo-EM. It takes a cryo-EM map and incorporates an external potential into an molecular dynamics (MD) simulation, guiding the atomic model to fit the experimental map while preserving physical realism. Several methods have been built on this idea, most commonly based on the correlation between an atomic structure and a reconstructed map [18–22] or 2D images [14]. Bayesian approaches shave been developed to match the average density from a simulated ensemble, rather than a single structure, to the observed map - effectively capturing the ensemble-averaged signal [**?**, 23, 24]. More recently, leveraging the advantages in automatic differentiation, machine learning methods have begun incorporating physical constraints - such as diffusion models - as priors to regularize atomic structures based on volumes or ensemble-averaged cryo-EM observables [12, 15, 17, 18, 25–28].

While these methods yield informative atomic models, they are not designed to recover the underlying ensemble distribution encoded in the cryo-EM particles—a feature essential for understanding biomolecular function—as they typically rely on fitting to averaged density maps constructed from subsets of the data. Recovering this ensemble distribution remains challenging, as the high noise levels in cryo-EM images render direct particle-to-class assignments unreliable [29]. Consequently, several methods have been developed to recover ensemble distributions through Bayesian inference or noise deconvolution [5, 30–32]. Some of these approaches require a pre-existing structural ensemble as input. For example, the ensemble reweighting method [33] depends on a prior ensemble that encompasses all relevant structural changes observed in the images, thereby limiting its applicability when the prior distribution does not fully cover the conformational space.

To address these limitations, we extend the ensemble reweighting framework with a physics-guided strategy that simultaneously optimizes atomic structures and their weights. Our iterative approach alternates between ensemble reweighting, unconstrained optimization of atomic positions based on a likelihood function derived from the cryo-EM images, and a physics-constrained regularization step that enforces structural plausibility under the physical model (e.g., the MD force field). By decoupling the cryo-EM optimization from the physics-constrained step, we reduce computational cost and enable more flexible analysis pipelines. The paper is organized as follows. We first provide background on cryo-EM ensemble reweighting, define the likelihood that relates cryo-EM images to atomic structures through a forward model, and introduce a structural prior to impose physical constraints. We then present the central theory of our work—the ensemble optimization method—followed by results on four benchmark systems of increasing complexity, with additional details provided in the Methods section at the end. We conclude with a discussion of our findings and outline directions for future work. We foresee that *cryo-EM ensemble optimization* will enable the inference of physically meaningful ensemble distributions from single-particle cryo-EM images of dynamic biomolecules.

## 2 Background

### 2.1 Single-particle cryo-EM

Cryo-EM experiments capture noisy 2D projection images of individual biomolecules flash-frozen in solution. Each particle image has an unknown projection direction and unknown conformation, represented by the positions of atoms 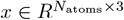, where *N*_atoms_ is the total number of atoms. If the system is at thermal equilibrium and the biomolecules are instantaneously frozen, each conformation is a sample from the Boltzmann distribution at the temperature prior to freezing. Knowing this distribution is essential, as it provides insight into the biomolecule’s thermodynamic properties, enabling us to study metastable states and understand the conformational transitions between them. In principle, we can infer this distribution from the cryo-EM images. However, this remains challenging due to the large number of unknowns and the high noise levels inherent in individual particle images.

### 2.2 Cryo-EM posterior probability for structural ensembles

The goal of the cryo-EM ensemble reweighting method [33] is to recover the biomolecule’s conformational probability distribution using individual cryo-EM images and a prior conformational ensemble, e.g., from MD or ML-based simulations. In this method, a family of probability densities *p*(*x* | *θ*) was proposed to approximate the Boltzmann distribution through Bayesian inference, where parameters *θ* reweight the prior ensemble. These are estimated by maximizing the posterior probability

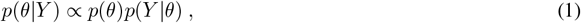

where 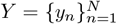 are independent and identically distributed (i.i.d.) experimental images, *p*(*Y* |*θ*) is the likelihood function, and *p*(*θ*) is the prior over parameters *θ*. The likelihood is the product over images of the image-to-structure likelihood, *p*(*y*|*x*) (section 2.3), marginalized over conformations

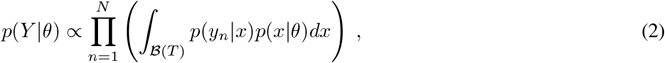

where ℬ (*T* ) is the domain of the Boltzmann distribution at temperature *T* .

To make the domain ℬ (*T* ) and distribution a concrete representation of the plausible molecular structures, the ensemble distribution *p*(*x*|*θ*) is parametrized through a discrete set of structures 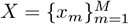 and their weights 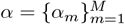 with *θ* = {*α, X*}

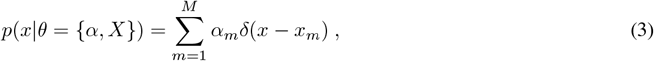

where *x*_*m*_ is an atomic structure representing a conformation and *α*_*m*_ corresponds to its weight. The weights are constrained to the simplex such that ∑_*m*_ *α*_*m*_ = 1 and *α*_*m*_ ≥ 0 ∀ *m*. In [5], the atomic structures *x*_*m*_ are considered known (e.g., found by clustering the MD prior), and only the weights *α*_*m*_ are optimized.

The primary contribution of this work is treating both *x*_*m*_ and *α*_*m*_ as unknown parameters. In previous work, the chosen atomic structures 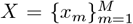 had physical meaning, but when we treat them as parameters without additional constraints and optimize over them, they quickly become physically implausible. Therefore, we set

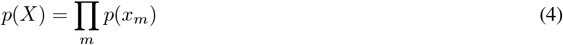

based on the physical likelihood of each individual structure (see section 2.4). Combining equations 2 and 3, and using Bayes’s theorem gives us the following posterior probability

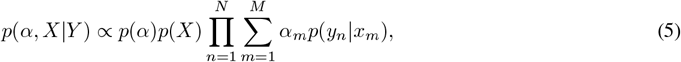

where *p*(*α*) is the prior of the weights.

### 2.3 Image-to-structure likelihood

The image formation process in cryo-EM connects the observed images 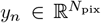 to an atomic structure 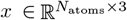. Assuming the weak-phase approximation [34, 35], a projection image *y*_*n*_ is generated by

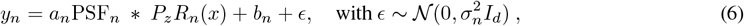

where * denotes convolution and *ϵ* is white Gaussian noise with variance 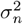. The parameters *a*_*n*_, *b*_*n*_ model a scale and a background offset factor in the experimental image [4]. The point spread function (PSF) operator models the imaging effects of the microscope, such as spherical aberration and astigmatism. The PSF is more commonly encountered in the literature by its counterpart in the Fourier domain, the contrast transfer function (CTF). For notation simplicity, we have omitted the explicit dependence of the PSF on the imaging parameters. The operator *P*_*z*_ projects the atomic structure along the z-axis, and *R*_*n*_ rotates and translates the molecule. The image-to-structure likelihood, *p*(*y* | *x*), is defined using this forward model by

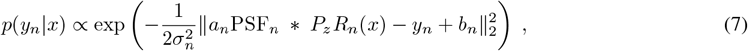

where 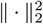 is the *l*^2^-norm. Additional details on this model and further considerations are described in the Methods section.

### 2.4 A physics-based prior

To construct the posterior probability defined by Eq. (5), we need to define the prior. In this paper, we construct our prior using an MD force field, an approximation to the Hamiltonian of biomolecules. Although other approaches are available to achieve this, such as diffusion models [25–27, 36] or structural samplers [37, 38], we defer their use to future work.

Let *E*_MD_ : 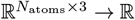 be the Hamiltonian defined by an MD force field. Then, the probability density function of the prior ensemble is given by

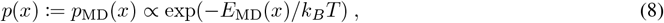

where *k*_*B*_ is Boltzmann’s constant and *T* is the temperature. MD force fields aim to approximate the physical model governing atomic interactions. The energy associated with these forces is designed to take on large values for physically unrealistic structures, such as those with steric clashes or broken bonds. While the energy takes continuous values, and does not provide a clear cutoff for what is “physical” vs. “not-physical,” the energy penalties for steric clashes and broken bonds are so significant that it is convenient for our discussion to view MD as restricting atomic structures to a domain of “physical” structures 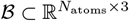.

## 3 The cryo-EM ensemble optimization method

We now build on the theory introduced in section 2 to develop the *cryo-EM ensemble optimization* method. The conceptual goal is to find the optimal set of structures 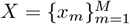 and corresponding weights 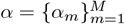 that maximize the posterior defined in Eq. (5), using the likelihood and prior from sections 2.3 and 2.4, respectively. The actual implementation is slightly more nuanced, primarily due to the following challenges: i) There is not one obvious approach to weighting the contribution of the MD energy and cryo-EM energy components, but it is enough for our purpose that the MD component excludes non-physical structures; ii) There are exceptional MD software packages that use elaborate models that include solvent interactions and temperature or pressure constraints in well-tuned simulations, which are difficult to integrate with other software directly or to reimplement; iii) MD simulations move molecules around in space, which would require us to track the viewing direction of each image with respect to all structures; iv) MD simulations are relatively expensive, and many expensive iterations may be required (depending on the way the components are combined).

To address these challenges, we adopt an iterative ensemble optimization approach conceptually inspired by projected gradient descent [41]. As illustrated in Figure 1, we decompose the overall optimization into four different steps. We fix the number of structures in the ensemble to *M* - a user-defined parameter. Inspired by the weighted ensemble and multiple walker metadynamics methods [42, 43], we refer to each member of the ensemble *m* = 1, …, *M* as a “walker”. The algorithm receives as input a dataset of cryo-EM images and their associated imaging parameters (CTF and pose information), a set of atomic structures, and a reference atomic structure to be used for alignment. Furthermore, optional inputs include an initial set of weights for the structures and a reference volume used to refine the alignment locally. Through these parameters the algorithm begins by generating an ensemble of initial structures (Initialization Step), and then iterates between structure alignment (step A), determining their weights based on the images (step B), optimization of the structures using only the cryo-EM likelihood (step C), and finally “projecting” the structures back onto the domain of the prior (step D). A more detailed overview of the steps is presented in the following sections; additional technical details are provided in the Methods section.

**Figure 1.**
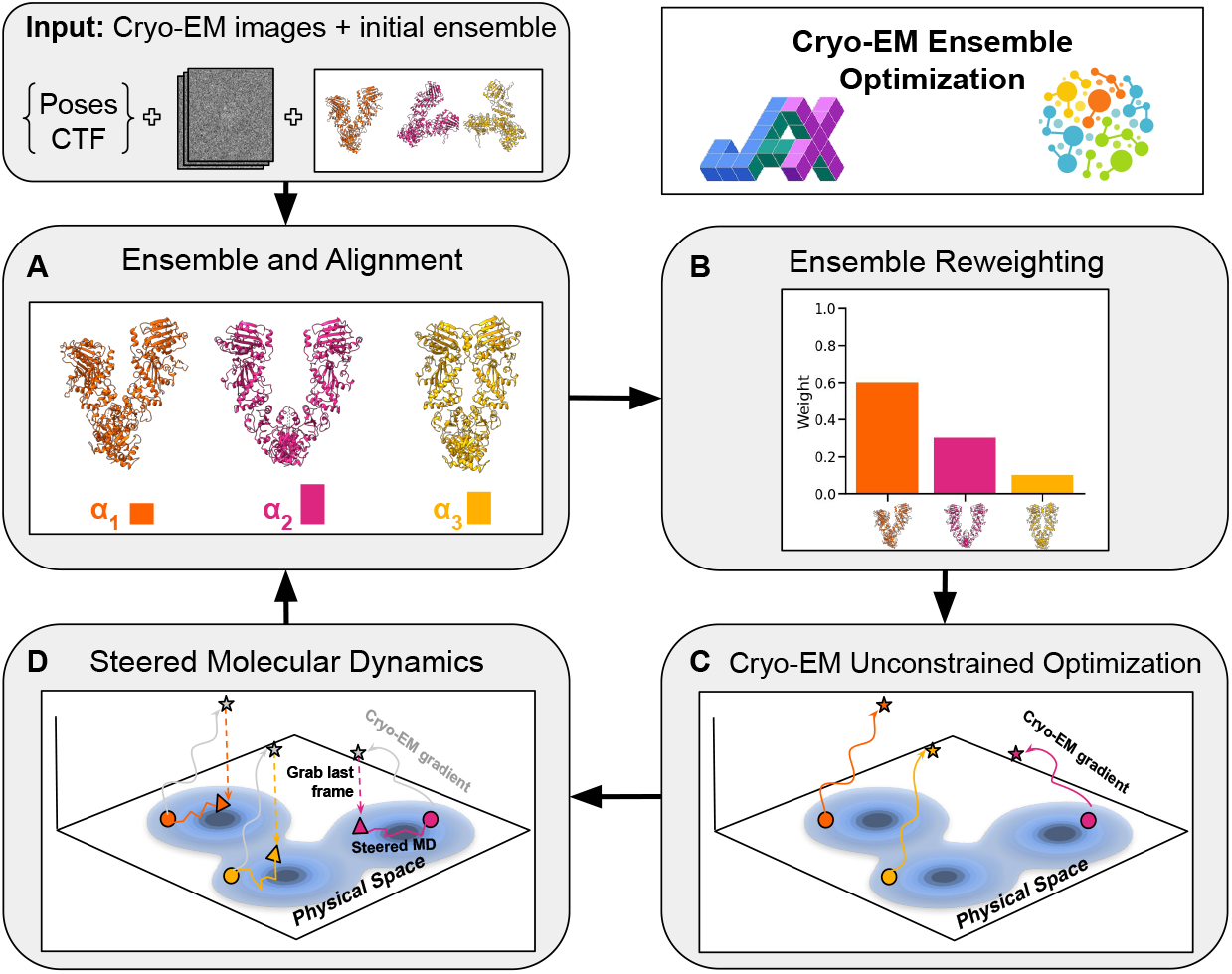
The cryo-EM ensemble optimization method. Starting from a structural ensemble (atomic models and weights) and cryo-EM particle images with imaging parameters (top left panel), we iterate over four steps. Step **A**: Define the ensemble and align the structures to a reference atomic model or map. Step **B**: Obtain the optimal weights for fixed structures by performing ensemble reweighting. Step **C**: With the weights fixed, perform unconstrained optimization on the atomic positions with the cryo-EM likelihood (Eq. (10)) as the target function. Step **D**: Project the optimized structures from Step **C** onto the domain of the physical prior by running steered MD. **Iteration:** Take the last frame from **D** as the structures and weights from **B** to define a new structural ensemble, and iterate until convergence is reached. The software is implemented using JAX [39] and OpenMM [40] (top right panel).

Our method is implemented as a module of the open-source library cryojax [44] and can be accessed at https://github.com/flatironinstitute/cryojax-ensemble-optimization.

### 3.1 Step A: structural ensemble and alignment

#### 3.1.1 Defining a structural ensemble

We define a structural ensemble as a set of structures 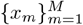 and their corresponding weights 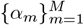. Before the first iteration, we can initialize the structures by running a short MD simulation from a reference structure, and selecting *M* random frames. Unless otherwise specified by the user, the initial weights are set to be uniform: *α*_*m*_ = 1*/M* . For following iterations, the structural ensemble is defined by the output of step B (weights) and step D (structures) outlined below.

#### 3.1.2 Structural alignment

To ensure an accurate comparison between the atomic structures and the experimental images, the structures *x*_*m*_ must be oriented so that the image computed using the forward model defined by Eq. (6) best matches the corresponding experimental image. Although we assume we know the optimal orientation for each particle, we need to align the structures to the frame of reference that defines this orientation. For the first iteration, we align the initial atomic structures with the map or reference structure that matches the experimental cryo-EM images (e.g., the consensus reconstruction volume). In subsequent iterations, the process is divided into two steps: a global alignment step and a local refinement step. For the global alignment, we use the Kabsch-Umeyama algorithm [45, 46] to align each walker to a pre-aligned atomic structure. This initial alignment is then locally refined using the reconstructed volume through rigid-body alignment. We use this two-step alignment process for two reasons: i) the initial structure might have a low correlation to the consensus volume, and thus the global alignment step might result in a structure that is not aligned to the image’s frame of reference, and ii) alignment to volumes is known to get stuck on local minima [47], which could result in the local alignment step producing a poorly aligned structure if the initialization is not close enough to the reference volume. Details of these procedures are provided in the Methods section. We note that this step could be replaced by a direct pose-refinement search, which we defer for future work.

### 3.2 Step B: Optimal weights for a fixed set of structures

With the aligned structural ensemble from step A, the second step consists of finding the weights that maximize the likelihood term in Eq. (5) (Figure 1B) for fixed *X*. We update *α* according to

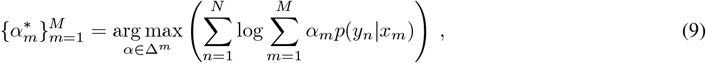

where Δ^*m*^ is the probability simplex of dimension *m*, with constraints 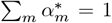 and 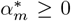. The image-to-structure likelihood, *p*(*y*_*n*_ | *x*_*m*_), is defined in Eq. (7). The solution can be obtained through expectation maximization or projected gradient descent on the probability simplex [48–50]. We optimize the log-likelihood for numerical stability.

### 3.3 Step C: Unconstrained ensemble optimization using the cryo-EM likelihoood

The third step involves optimizing the structural walkers by maximizing the likelihood term in Eq. (5), starting from the structures obtained in step A, and utilizing the optimal weights from step B (Figure 1C). Since cryo-EM datasets are usually large, we approximate this solution by running stochastic steepest descent with uniformly random subsets of the images (*Y*_*s*_ ⊂ *Y* ). We calculate the likelihood for a given subset as

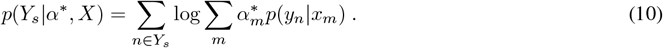

We use the steepest descent method because it conveniently allows us to specify the step size in Angstroms, making this parameter more relatable to atomic structures. The number of optimization steps and the step size are user-defined. To compute the gradients, we utilize automatic differentiation through JAX [39] via the differentiable simulator implemented in cryoJAX [44]. Using automatic differentiation allows us to implement more complex forward models while keeping the implementation relatively simple.

It is important to note that the structures obtained through this optimization are unlikely to lie within ℬ, the domain of physical structures, since the optimization is unconstrained with respect to the physical and chemical properties of the biomolecule. We denote the output of this step as 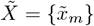, an intermediate collection of possibly non-physical walkers that is corrected in the next step.

### 3.4 Step D: projecting onto the physical prior

The fourth step (Figure 1D) involves regularizing the output from Step C by “projecting” it onto the domain ℬ of physical structures, a heuristic inspired by projected gradient descent [41]. For each structure 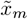 obtained in step C, we aim to find

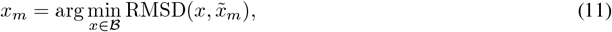

where RMSD is the root mean square deviation: 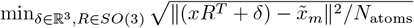, where *R* and *δ* are a rotation and a translation, respectively. The minimization problem defined by Eq. (11) does not have an analytic solution, and, as we pointed out, the domain ℬ is not formally defined. Instead, we find an approximate solution by running a steered MD simulation for each walker *m*, with a force field defined as

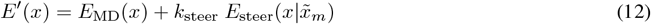

where 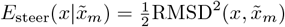 is the steering energy, and *k*_steer_ is the steering constant. The steering energy indirectly incorporates information from the cryo-EM data into the MD simulation. The steering constant *k*_steer_ balances the contributions of the prior and the cryo-EM likelihood. Selection of an appropriate value is described in section 6.2.4. The steered MD simulation is run for a user-defined number of steps, where the last frame of each simulation is used to define a new set of structures. This step is analogous to the projection step in projected gradient descent. Specifically, for each walker, we identify the closest physical structure to 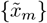 that satisfies the constraints imposed by the force field. Note that the biasing constant, *k*_steer_, could be alternatively added to *E*_MD_, effectively changing the temperature of the system.

In traditional cryo-EM integrative modeling approaches, such as MDFF [17], the cryo-EM “energy” term is computed by directly comparing simulated atomic structures with the cryo-EM reconstruction. We deviate from this approach by computing *E*_steer_ using a surrogate function, namely the RMSD, which enables an indirect comparison between simulated atomic structures and cryo-EM images. This design choice offers three key advantages: (i) direct comparison between atomic models and cryo-EM images across large datasets is computationally and memory-intensive; (ii) high-frequency variations—such as those arising from individual MD steps—are often obscured by noise and the CTF, contributing minimally to *E*_steer_, while stable MD simulations requiring small time steps ( ∼ 1 fs) necessitate millions of steps, making direct computation at each step impractical; and (iii) the coupling between walkers in the likelihood would require running multiple simulations in parallel, substantially increasing computational demands and complicating implementation.

### 3.5 Iteration and convergence

The cryo-EM ensemble optimization method consists of iterating steps A-D (Figure 1): defining and aligning the structural ensemble (A), the weight optimization (B), unconstrained optimization with the cryo-EM likelihood as the objective function (C), and a projection-like step through steered MD (D), as described above. More details are provided in the Methods. In the following examples, we assess method convergence by monitoring the behavior of cryo-EM likelihood. When ground-truth ensembles are available, we also monitor the RMSD and the weights relative to these reference structures as a function of the iteration step. A systematic criterion for convergence is left to future work.

## 4 Results

We evaluate the performance of cryo-EM ensemble optimization on several benchmark systems of increasing complexity. Our goal is to assess whether the method can recover the optimal structures and weights across different prior landscapes and under diverse, realistic cryo-EM imaging scenarios.

### 4.1 The rectangle: a simple toy system

We begin by evaluating our method on a simple 2D toy system of four typeless atoms that form a rectangle in the *xy*-plane. Although far from a real biological specimen, we chose this toy model as it allows us to easily design different priors and, thus, test our method against different physical conditions. In addition, visualization and data generation for this system are simple, as any configuration can be identified by two intuitive collective variables: the height (*h*) and length (*l*) of the rectangle. We design the physical prior of the rectangle such that only rectangular configurations are possible, which we achieve by including an energy term that depends on the angles of any pair of 3 atoms. Additionally, to closely mimic biological force fields, we define wells and barriers, represented by a harmonic potential, at arbitrary configurations, effectively making some rectangular configurations more or less likely in the prior distribution. To simulate data, we sample heights and lengths from a 2-dimensional multimodal distribution, use these values to define rectangles, and simulate one image from each sample using the imaging forward model defined in Eq. (6). We define the heights and lengths distribution as a weighted set of three Gaussians, where the weights determines the number of samples obtained from each Gaussian. We refer to each mode in the distribution as a metastable state, and identify them as *S*_*i*_ for *i* = 0, 1, 2, with representative samples shown in Figure 2A. For our experiments, we set the weights to (0.125, 0.250, 0.625) and simulate a total of 800 images. Example images are shown in Figure 2A, identified by the height and lengths used to generate them. Details on the definition of the physical prior are provided in the Methods section.

**Figure 2.**
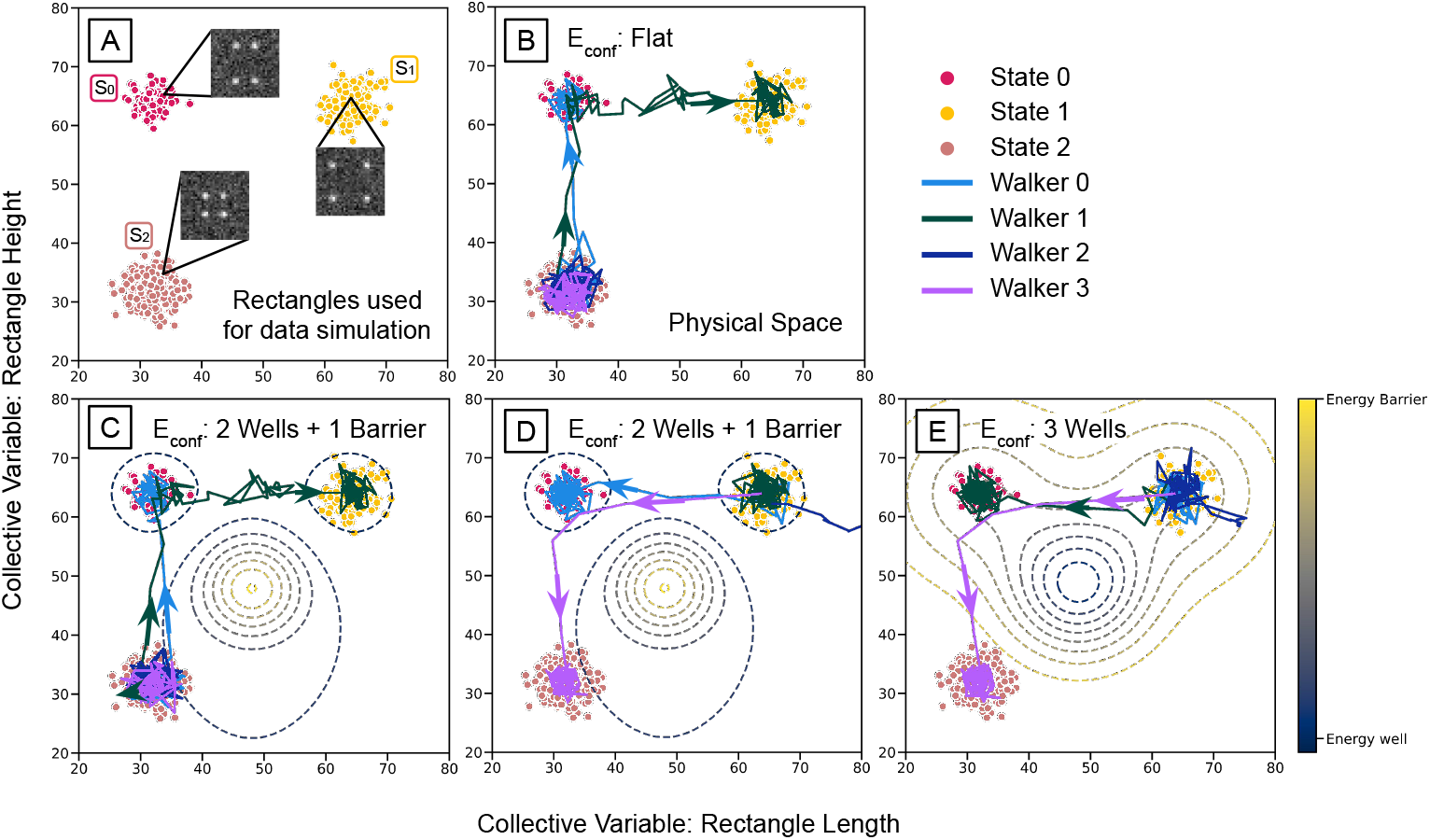
Cryo-EM ensemble optimization results for a rectangular toy model using different physical priors and initialization schemes. (A) Distribution of rectangular conformations used to generate synthetic cryo-EM images, characterized by their height and length. There are three metastable states, each consisting of 800 images labeled *S*_*i*_ with weights (0.125, 0.250, 0.625), respectively. Example images are shown for each state. (B) Ensemble optimization trajectory for four walkers with a physical prior where all rectangles are equally likely. The initial structures are located near the center of the largest metastable state, *S*_2_. (C) Ensemble optimization results for the same cryo-EM image distribution, incorporating a multimodal prior in the rectangular conformational space, which includes two energy wells and one energy barrier. The centers of the energy wells correspond to the centers of the image states *S*_0_ and *S*_1_, and the energy barrier lies at the center of the landscape. The walkers are initialized as in (B). (D) The same prior as in (C) is used, but the walkers are initialized close to *S*_1_. (E) The energy barrier from (C) is switched to an energy well deeper than the other wells, and the method is initialized from *S*_1_. In all cases, the arrows show the direction of each walker’s trajectory.

Figure 2B-E shows examples of the method’s performance with different priors and initializations. First, we consider the case where all rectangles are distributed uniformly in the prior (Figure 2B). We initialize our method with four walkers (*M* = 4), starting from the most populated metastable state (*S*_2_) and assigning equal initial weights to each walker. We run 100 iterations of the ensemble optimization procedure. The walker trajectories are shown as lines with arrows. With this setup, we show that our method can explore less-populated states in the distribution of cryo-EM images, even when initialized near the most populated state, while maintaining rectangular geometries (see Supporting Information). This can be observed in the trajectories of walkers 1 and 2 (green and light blue, respectively; Figure 2B), which escape *S*_2_ and reach the other metastable states. Once walker 1 arrives at *S*_0_, walker 2 changes its trajectory and aims for *S*_1_. Meanwhile, the remaining walkers explore different conformations of *S*_2_. The estimated weights for each walker are (*α*_0_, *α*_1_, *α*_2_, *α*_3_) = (0.125, 0.250, 0.224, 0.401). If we assign walkers to metastable states such that walkers 2 and 3 correspond to *S*_2_, the estimated state weights are (0.125, 0.250, 0.625), which match the true weights.

Now, we focus on cases where the prior conformational distribution is not flat. We add terms to the force field used to run the MD: two high-probability states (energy wells) are added to *S*_0_ and *S*_1_, and a low-probability state (energy barrier) is placed in the middle of the landscape. We show the results for two instances of this test in Figure 2C-D, with initialization being the only difference. In Figure 2C, we start, as in the previous test, from *S*_2_, while in Figure 2D, we start from *S*_1_ (the lowest populated state). We find that the walkers overcome the barrier, as demonstrated by their transitions to different conformations and by their escape from local minima in both image and physical space. Additionally, energy barriers affect how the walkers move through the optimization process. In Figure 2D, one of the walkers drifts outside of the metastable state, and is assigned a weight of 0. As in Figure 2B, the optimized weights for each metastable state match the actual values.

In our final test for the rectangle toy system, we replace the barrier with an energy well deeper than those at *S*_0_ and *S*_1_. We start from rectangles from *S*_1_. In Figure 2E, we show the trajectories of each of the walkers. During the optimization, the walkers that transition to other metastable states do so without drifting towards the deep energy. This test shows that our method allows walkers to escape local energy minima in the prior that are not represented by the images. The estimated state weights closely match the actual values.

### 4.2 Alanine tripeptide

We proceed to study a slightly more complex system, alanine tripeptide, a commonly used toy system in MD [42, 51, 52]. We use this system to highlight the role of physical constraints in the optimization process. Specifically, we compare the full pipeline with a variant that omits the projection step onto physical space, i.e., optimizing the structural ensemble using only the cryo-EM images. This comparison illustrates the importance of incorporating a physical prior.

For this test, we simulated images from two distinct conformations, denoted 3Ala-A and 3Ala-B, with probabilities 0.7 and 0.3, respectively (details in section 6.4.2). Figure 3 presents the results of running the pipeline with and without the projection onto the physical space in Step D. For each case, we use two walkers and initialize the algorithm with both walkers starting from a third conformation, 3Ala-C. In the unconstrained case (without a prior), the final optimized structures exhibit broken bonds due to the absence of physical constraints (Figure 3C). Although the cryo-EM likelihood (Eq. 17) improves, the heavy-atom RMSD relative to the true structures shows minimal improvement, demonstrating that without a physical prior, the method overfits to non-structural information in the images (Figure 3A). In contrast, incorporating a physical prior yields physically reasonable optimized structures without broken bonds (Figure 3C), marked improvement in RMSD relative to the true structures (Figure 3B), and superior likelihood compared to the unconstrained case (Figure 3A). These results show the importance of physical constraints, not only for avoiding bond breakage but also for preventing overfitting due to spurious features in the images, such as noise. The ensemble weight for the walker closest to 3Ala-A was 0.70 in the unconstrained case and 0.74 with the physical prior.

**Figure 3.**
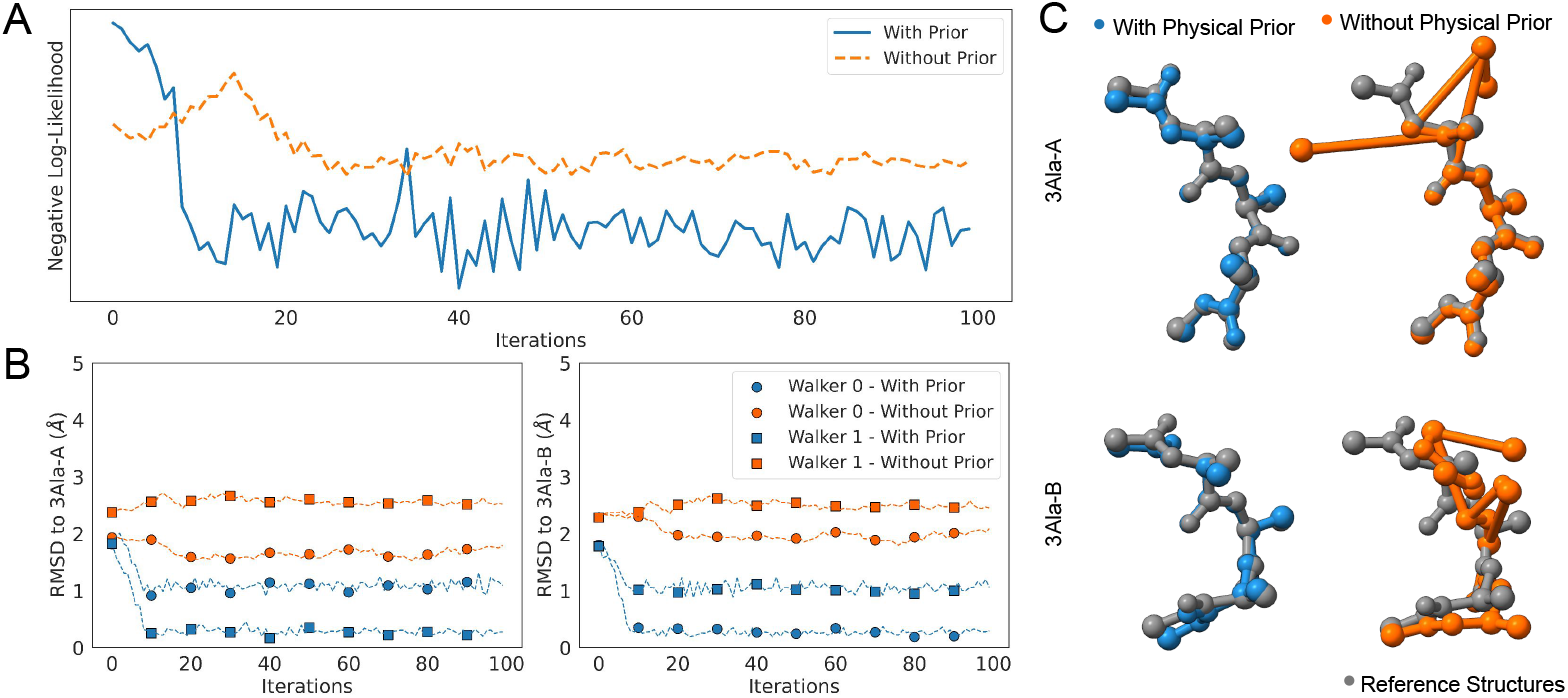
Cryo-EM ensemble optimization results for tri-alanine with and without the physical prior. For the case without a prior, we omit Step D in Figure 1. We generate 5000 synthetic images from two conformations, 3Ala-A and 3Ala-B, shown at the top and bottom of panel C, respectively, with 70% of the images generated from 3Ala-A. (A) Negative log-likelihood (negative of Eq. (17)) as a function of iteration for the ensembles estimated with prior (blue, solid) and without prior (orange, dashed). (B) Heavy-atom RMSD relative to the true atomic models (left: 3Ala-A; right: 3Ala-B) for walkers evolved using both approaches. (C) Optimized structures with a physical prior (blue, left), and without a physical prior (orange, right). The results demonstrate that using a physical prior yields structures that more closely match the references and the experimental data, indicating that the absence of a prior can lead to broken bonds and result in being trapped in local minima.

### 4.3 Chain A of GroEL

In this section, we benchmark our method on the chaperonin GroEL, focusing on how the optimized ensemble is affected by spurious artifacts in the experimental data, such as increased imaging noise or uncertainties in the imaging parameters. Our tests are inspired by [23], where the authors construct a heterogeneous ensemble by extracting chain A of the apo conformation of GroEL, and building a homology model for the same chain of GroEL-ADP in the relaxed allosteric state. We refer to these two states as apo-GroEL and holo-GroEL, respectively. We conduct two tests: first, evaluating our method on images generated at different signal-to-noise ratios (SNRs), and second, examining how uncertainties in particle poses affect the final ensemble. Additional details about the setup are provided in section 6.4.3.

In the first test, we generate a homogeneous particle set and examine how closely the pipeline can resolve a walker to the true atomic model. Specifically, we simulate 1,000 images from holo-GroEL and run our method with a single walker initialized from apo-GroEL, assigned 100% of the weight. We repeat this test across multiple simulated datasets, each generated at a different SNR. We compare the results to the ideal case, defined as the scenario in which the true structure is known and the solution can be obtained by performing a steered MD toward it (i.e., executing only Step D in Figure 1 using holo-GroEL as the reference). This represents the lowest RMSD achievable relative to holo-GroEL with the steered MD setup. Figure 4A shows results at SNR values of 10^1^, 10^0^, 10^−1^, and 10^−2^. For all the SNR values tested, the walker converges to a structure with a heavy-atom RMSD to holo-GroEL within 1.7 − 1.9 Å, with the value increasing inversely proportional to the SNR. In all cases, the algorithm converges in fewer than 20 iterations, corresponding to an aggregated MD simulation time of 0.02 ns. The ideal case achieves a 1.35 Å heavy-atom RMSD to holo-GroEL. These results demonstrate that cryo-EM ensemble optimization yields structures that closely resemble those observed in the images, achieving convergence within a relatively short MD simulation time.

**Figure 4.**
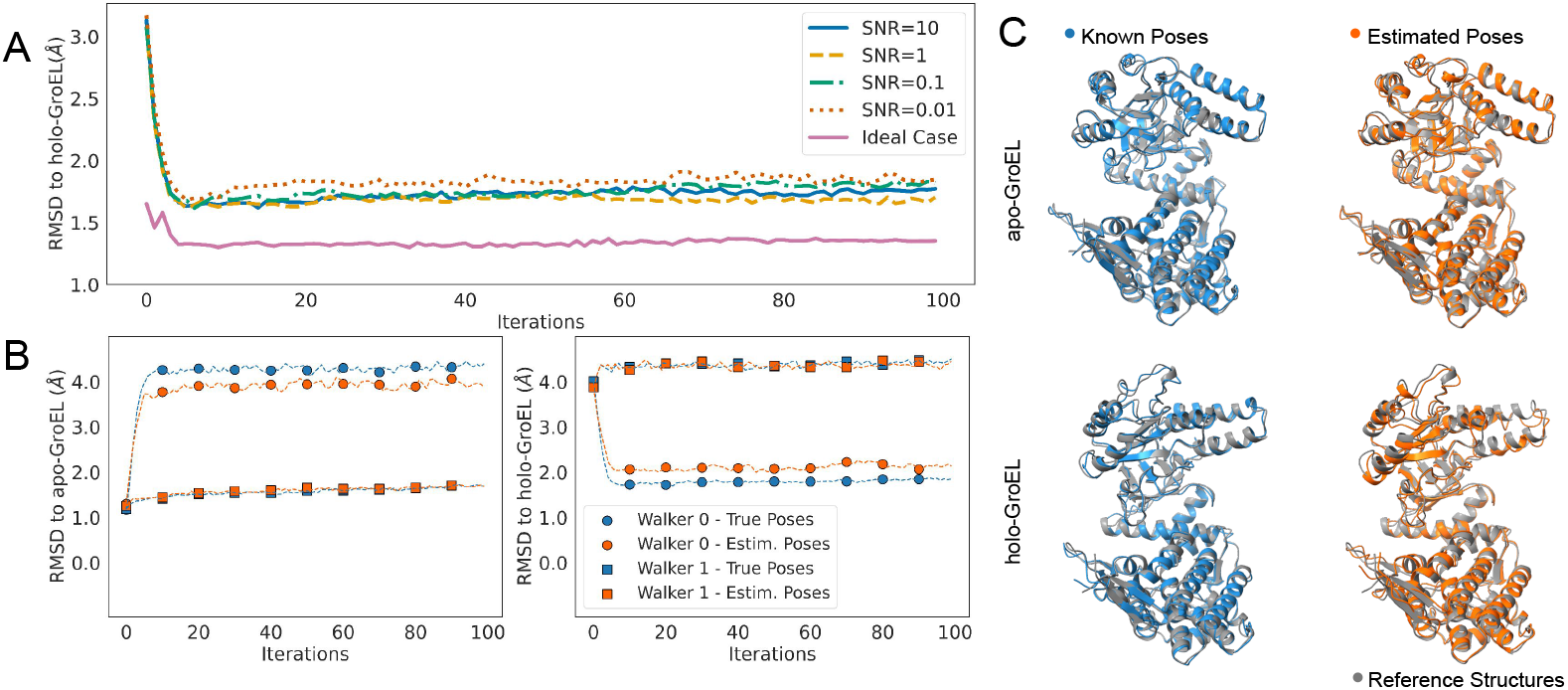
Cryo-EM ensemble optimization results for GroEL under varying noise levels and pose uncertainties, using an MD force field as a physical prior. (A) Ensemble optimization results at different noise levels using 1,000 images each generated from holo-GroEL. The method begins with a single walker initialized from the apo-GroEL state. The heavy-atom RMSD to holo-GroEL is shown over iterations, with the ideal case (steered MD using holo-GroEL as a reference) indicated by a pink solid line. (B) Two instances of ensemble optimization are compared: one using ground-truth poses and the other using estimated poses. The left and right subplots display the heavy-atom RMSD to apo-GroEL and holo-GroEL, respectively. Walkers with known poses are shown in blue, and those with estimated poses are shown in orange. (C) Apo-GroEL and holo-GroEL optimized conformations (blue/right and orange/left, respectively). The reference structures for the different cases are shown in gray.

The second test focuses on how uncertainties in the pose parameters affect the optimized cryo-EM ensemble. Usual cryo-EM heterogeneity methods assume that the poses are known — typically obtained through an *ab initio* reconstruction process — which can lead to inaccuracies in the poses. For this test, we simulate a heterogeneous dataset by generating 20000 images from apo-GroEL and holo-GroEL. To create each image, we sample one of the two conformations with probabilities 0.7 and 0.3, respectively, having an SNR of 0.1 (see section 6.4.3 for more details). We estimate the pose parameters by performing *ab initio* homogeneous reconstruction and refinement in CryoSPARC [53] using the entire image set. Note that since the homogeneous volume represents an average of the conformations in the dataset, the *ab initio* poses may be slightly inaccurate. We run two instances of the test, one using the ground-truth poses and the other using the *ab initio* poses. In each instance, we initialize two walkers from apo-GroEL and run the complete cryo-EM ensemble optimization. Figure 4B shows the heavy-atom RMSD to each reference structure as a function of iteration step. Using the true poses slightly improves recovery of the holo structure, reducing the heavy-atom RMSD to holo-GroEL by 0.2Å compared to the structure optimized using estimated poses. Both cases result in an accurate estimation of the weights, using known poses yields values (*α*_*apo*_, *α*_*holo*_) = (0.70, 0.30), whereas estimating the poses leads to a slight underestimation of the holo-structure weight (*α*_*apo*_, *α*_*holo*_) = (0.71, 0.29).

### 4.4 Real data of Hsp90

We apply the cryo-EM ensemble optimization pipeline to the hsp90-p23-GR dataset (EMPIAR-11028). Since the true ensemble distribution is unknown for real data, we test the optimization on a single subset of real particles that reconstruct to high resolution—specifically, those assigned to hsp90–p23. [54]. To challenge the method, we initialize with a distant conformation (heavy-atom RMSD to the deposited structure of 10Å). Furthermore, we remove the p23 protein to test the robustness of our method against model misspecification. The initial structure is obtained from the atomic models designed in [55], and it is illustrated in Figure 5A. We downsample the images to a size of 128 × 128 (Figure 5A). We use the CTF parameters estimated in [54], and re-estimate the pose parameters through *ab-initio* reconstruction. We run the method using the images corresponding to the first half-map, referred to as the *training* dataset, and those assigned to the second half-map are left for cross-validation - an independent set not used to optimize the structures- and are referred to as the *test* dataset. We initialize the method from the conformation defined above and run the pipeline with a single walker assigned 100 % weight for a total of 100 iterations. Figure 1B compares the initial and optimized structures, demonstrating the ability of our method to induce large conformational changes. Furthermore, Figure 5B shows that the negative log-likelihoods systematically decrease for both the *training* and *test* sets. Variability across the test sets was quantified using a 95% confidence interval over 1000 independent subsets. The behaviour of the negative log-likelihood across the test data indicates that the optimized structure is not overfitting to the training data. Similarly, Figure 5C shows that the final optimized structure accurately models the consensus volume from [54], as evidenced by high local correlation and clear improvement in the Fourier shell correlation (FSC) area under the curve.

**Figure 5.**
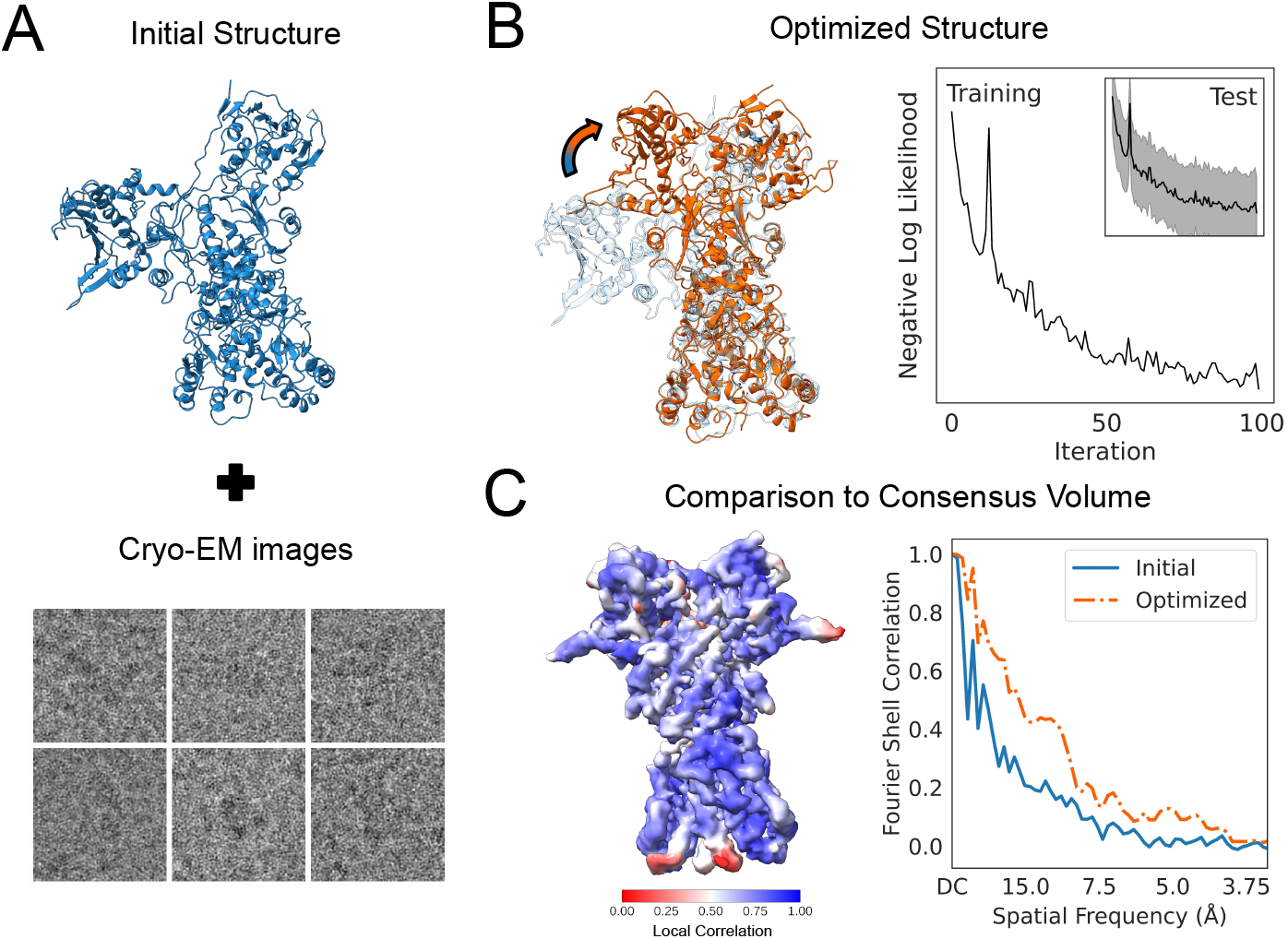
Cryo-EM ensemble optimization results for the hsp90-p23 particle subset of EMPIAR-11028. (A) Structure used to initialize the ensemble optimization method (blue) and representative images from the dataset. (B) Left: optimized structure (orange) superimposed with the initial structure (blue), showing the conformational changes induced by optimization. Right: negative log-likelihood (negative of Eq. 17) for the training (main panel) and test particle sets (inset). For the inset, we compute the value of the loss for 1000 random subsets of the test dataset, and define the error bars using a 95% confidence interval. (C) Left: local correlation between the consensus volume and optimized structure. Right: Fourier shell correlation (FSC) between the consensus volume and maps generated from the optimized structure (orange dashed line) and the initial structure (solid blue line).

## 5 Conclusions and Discussion

In this work, we developed a method for optimizing ensemble distributions from single-particle cryo-EM images using physical structural priors. Our approach effectively balances physical information and experimental data to produce structurally feasible ensembles with their respective weights, thereby approximating the conformational distribution observed in the images. To evaluate method performance, we conducted benchmarks on increasingly complex systems: a simple toy model of four typeless atoms, alanine tripeptide, GroEL chain A, and Hsp90. The toy model provides insights into how the method responds to variations in the prior landscape, while the tri-alanine benchmark demonstrates the importance of physical constraints during ensemble optimization, extending beyond reliance solely on cryo-EM images. The GroEL system illustrates the impact of data inaccuracies—specifically, imaging noise and pose uncertainty—on method performance. Finally, the Hsp90 benchmark tests the method under challenging conditions: poor initialization, model misspecification, and real experimental data. Across all benchmarks, including cases with imperfect pose estimates, the method accurately infers metastable state conformations and populations. Notably, our approach does not require dimensionality reduction or collective variables, making it readily generalizable to complex, flexible systems.

Our method is built on cryoJAX [44], JAX [39], and Equinox [56], enabling automatic differentiation and efficient GPU computations. For MD simulations, we use OpenMM [40], a flexible MD engine with a Python API. This combination provides computational efficiency while maintaining Python’s flexibility. The implementation is sufficiently modular to integrate with other single-particle techniques, requiring only a likelihood-equivalent metric and forward model. Furthermore, the framework accommodates diverse physical priors, including alternative force fields and modeling approaches, such as diffusion models, that can provide unnormalized structural energy terms.

The cryo-EM ensemble optimization method presented here has several limitations. Like other cryo-EM heterogeneity methods, we assume poses are known. However, this assumption breaks down in high-noise or highly heterogeneous settings, where typical *ab initio* reconstruction pipelines yield less accurate pose estimates. Addressing this limitation would require an additional pose-optimization step, thereby increasing computational cost. While computational cost scales linearly with the number of walkers, MD simulation of large, flexible systems may become a bottleneck, necessitating more efficient simulation approaches such as coarse-grained MD or replacing the MD engine with a diffusion model. Nevertheless, this work demonstrates a viable path forward for inferring ensemble distributions of biologically relevant macromolecules directly from cryo-EM images, opening new avenues for understanding biomolecular dynamics and conformational heterogeneity.

## 6 Methods

In this section, we provide a detailed description of the imaging forward model, each step of the cryo-EM ensemble optimization method, and the specifics for the benchmark systems.

### 6.1 Cryo-EM imaging forward model

The cryo-EM imaging forward model (Eq. 6) transforms structure through the following steps: rotation and translation, projection along the *z*-axis, and application of microscope imaging effects via the PSF. Details for each step, including the full image-to-structure likelihood, are provided below.

#### 6.1.1 Pose

The pose parameter specifies the orientation and position of the structure in the image. We assume that the pose of each image is known and represent it using a rotation and translation operator *R*_*n*_. These parameters can be obtained from a consensus reconstruction process, and we leave their optimization for future work. It is important to consider that the structure *x*_*m*_ must be aligned to a reference to which the poses are calculated, e.g., the volume obtained via the consensus reconstruction process. The atomic structures are rotated and translated before the projection operator is applied, as described below.

#### 6.1.2 Projection operator

The projection operator takes a rotated and translated atomic structure *x* and makes a noise-free image of it by projecting its density along the *z* axis. We define the projection operator in Eq. (6) using the atomic density parameterization defined in [57, 58], where each atom is assigned five Gaussians with the center being the atomic coordinate, and the amplitude and variance defined by the atomic electron form factors. The projection, 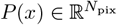, is discretized to a 2D grid with *N*_pix_ pixels. Therefore, given a structure 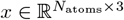, the clean 2D projection is given by

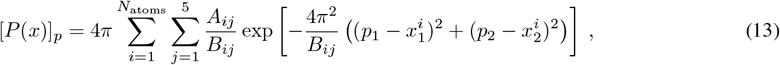

where *p* = (*p*_1_, *p*_2_) denotes the pixel coordinates in the discretized 2D grid, and *A*_*ij*_ and *B*_*ij*_ are the electron form factors. The *i*-th atom of the structure, *x*^*i*^, is represented as a 3D vector with entries 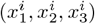. When available, the Debye-Waller factors are added to the *B*_*ij*_ electron form factors to account for the temperature-induced vibration of the atoms [59], that is, we redefine *B*_*ij*_ → *B*_*ij*_ + *b*_*DWi*_, where *b*_*DWi*_ is the Debye-Waller factor for atom *i*. We obtain the electron form factors using the Python library gemmi [60], and the atomic structures and their b-factors can be obtained from the Protein Data Bank [61].

#### 6.1.3 Point spread function

We define the point spread function (PSF) in terms of its Fourier-domain counterpart, the Contrast Transfer Function (CTF). We use the anisotropic CTF defined in CTFFIND4 [62] and implemented in cryoJAX [44]. The CTF parameters are assumed to be known and can be obtained through the typical cryo-EM single particle analysis pipeline. The CTF is applied in the Fourier domain following the Fourier Convolution Theorem.

#### 6.1.4 Image-to structure likelihood

The likelihood *p*(*y*_*n*_ | *x*_*m*_), Eq. (7), compares each image *y*_*n*_ to structure *x*_*m*_. To estimate it, we must account for the rotation and projection of the structure, convolution with the PSF (as described above), the noise parameters, and the scale and offset factors. We estimate the scaling *a*_*n*_ and offset *b*_*n*_ factors on-the-fly for each image, by solving the following minimization problem

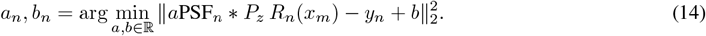

Details on the solution of this optimization problem can be found in the Supporting Text A.1.1. We marginalize the likelihood *p*(*y*_*n*_|*x*_*m*_) over the noise variance *σ*^2^. We assume a uniform prior for the variance, resulting in

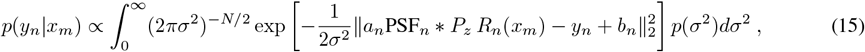

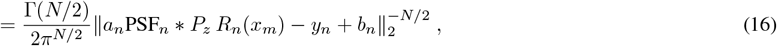

where we defined the prior for the variance using the Jeffrey’s prior [63, 64] for the variance of a multivariate isotropic Gaussian distribution, *p*(*σ*^2^) ∝ 1*/σ*^2^. Details of the derivation can be found in the Supporting Text A.1.2.

#### 6.1.5 Ensemble likelihood

The ensemble-to-images likelihood can be obtained directly from the posterior probability defined in Eq. (5), expressed as

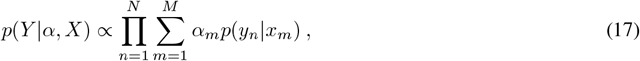

where *p*(*y*_*n*_|*x*_*m*_) is the image-to-structure likelihood defined in Eq. (16).

### 6.2 Cryo-EM Ensemble Optimization

The cryo-EM ensemble optimization method is an iterative optimization scheme inspired by the projected gradient descent algorithm [41]. Generally, an iteration of the method involves aligning the walkers to the images’ frame of reference (step A), reweighting the walkers (step B), performing unconstrained optimization on the walkers’ positions given a set of cryo-EM images (step C), and projecting the optimized walkers back to physical space using a physics-based approach (step D). Further details of each step are provided below.

#### 6.2.1 Step A: Defining a structural ensemble and alignment

##### Initializiation

The structural ensemble is initialized with *M* atomic models, where *M* is chosen by the user. The collection of atomic models at each iteration is referred to as walkers, denoted by 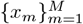. The ensemble distribution is defined by the set of walkers together with their associated weights. The initial walkers can be generated either by running a short MD simulation starting from a single structure and selecting random frames, or by using distinct conformations directly. Details on the initialization procedure for each system considered in this work are provided below. The initial weights can be defined by the user and are considered uniform by default. For subsequent iterations, the structural ensemble is derived from the output of the projection step D and its weights from step B.

##### Structural Alignment

The user must provide an atomic structure that has been pre-aligned with the frame of reference of the images. This can be achieved, for example, by aligning the initial structure to the cryo-EM consensus volume. In our benchmarks, we perform this alignment manually with UCSF ChimeraX [65]. Furthermore, the consensus volume can be provided as an optional input for realigning atomic structures at each iteration of the method.

In steps B and C, we compare the walkers to the images via the image-to-structure likelihood defined by Eq. (7). This requires evaluating the forward model described in section 6.1, which involves a rotation and a translation. To ensure a proper comparison between the images and the structures, the structures must be oriented so that evaluating the forward model with a given set of imaging parameters yields a simulated image that matches the corresponding experimental image.

We address the alignment issue by realigning the walkers in two steps. First, we align the walkers to a user-defined pre-aligned atomic structure using the Kabsch-Umeyama algorithm [45, 46]. This algorithm determines the rotation and translation that minimize the Euclidean distance between two atomic models, providing an initial estimate of the optimal rotation and translation. Second, we refine this alignment locally by rigidly aligning each walker with the reconstructed volume. While the local refinement step is susceptible to local minima that may yield suboptimal orientations, the initial global alignment helps mitigate this issue.

The rotation *R* ∈ *SO*(3) and translation *δ* ∈ R^3^ that solve rigid-body alignment between an atomic structure, 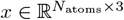, and a volume, *V*, are obtained by maximizing the correlation to the reference volume

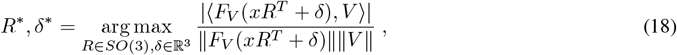

where *F*_*V*_ is a structure-to-volume forward model (see Supporting Text C), and | · |,∥ · ∥,⟨ · ⟩, denote the absolute value, *ℓ*2-norm, and the Euclidean inner product, respectively. We solve this optimization problem using gradient-based optimization with Optax [66] and the structure-to-volume forward model implemented in cryoJAX [44]. For simplicity, we optimize the rotation by representing it as a quaternion, which we constrain to lie on the unit 4-sphere after each optimization step.

#### 6.2.2 Step B: Optimal weights for a fixed set of structures

Using the aligned structural ensemble from Step A, we follow the ensemble reweighting method [33] to obtain the weight *α*_*m*_ of each walker *x*_*m*_. We assume that the *x*_*m*_ are fixed. We approximate the solution to Eq. (9) by running projected gradient descent on the simplex using JAXopt [67]. To improve computational performance, we perform this reweighting utilizing a subset of the images (see details below). Numerical and statistical reasons, the discussion of which we defer to a separate paper, may lead to weights close to zero, resulting in vanishing gradients. To prevent this, we apply a softmax function to the estimated weights 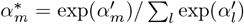 where 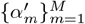 is the set of weights obtained through the projected gradient descent optimization. The softmax function helps us maintain the hierarchy of the walkers while avoiding vanishing gradients. This approach can lead to some weights being over- or under-estimated. Therefore, we re-estimate the weights using the entire dataset as a post-processing step, performed after the final iteration of the method.

#### 6.2.3 Step C: Unconstrained ensemble optimization using the cryo-EM likelihoood

In this step, we optimize the atomic positions for the walkers using the cryo-EM likelihood with a given subset of images, Eq. (10), denoted by *Y*_*s*_ ⊂ *Y* . We limited the optimization to a subset of atoms, indexed by *J*, which in our tests consists of all atoms except hydrogen. Then, we run steepest descent, with the update rule for each atom *i* defined as

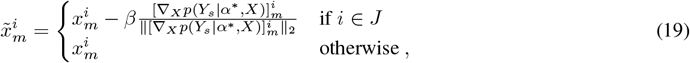

where *β*, the step size, is defined in Angstrom.

##### Image mini batching

As mentioned before, to avoid local maxima and improve computational performance, we employ a stochastic approach for running the optimization steps. Specifically, we sample a subset of the images, *Y*_*s*_ ⊂ *Y*, perform weight optimization, and then optimize the walkers’ atomic positions (steps B and C, respectively). We iterate *T* times before moving to Step D, with *T* being a hyperparameter.

#### 6.2.4 Step D: Projecting onto the physical prior

Since the optimization in the previous step is unconstrained, the optimized atomic structures are unlikely to satisfy the physical constraints imposed by the prior. With this step, we aim to find structures that satisfy the physical constraints and are as close as possible to the output from step C in RMSD. We achieve this by running a steered MD simulation for each walker, with the energy term defined in Eq. (12), where the optimized structures are used as the reference structure in the steering force. Furthermore, only the subset of atoms used for the optimization (those indexed by *J* in Eq. (19)) is used when computing the steering energy term *E*_steer_.

A key aspect when running the optimization step is properly weighting the contribution of the two energy terms, *E*_MD_ and *E*_steer_, which we control through the steering constant *k*_steer_ (see section 3.4). We choose *k*_steer_ such that the steering force (*F*_steer_) is a user-defined percentage *γ* of the total force (*F*_MD_ + *k*_steer_*F*_steer_). To do this, we run a short, unbiased MD simulation. For each of these time frames, *t*, we solve for the bias constant *k*_steer_ given the desired percentage (*γ*):

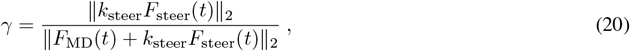

where the forces are related to the energies as: *F*_steer_(*t*) = − ∇ *E*_steer_(*t*) and *F*_MD_(*t*) = − ∇ *E*_MD_(*t*); and the biasing force is computed with respect to the initial positions of the simulation. In addition, the norms of the forces are computed using only the forces applied to the atoms whose positions are optimized in Step C. The final steering constant is set to the average over all time frames from the unbiased simulation. A detailed derivation of this quantity is provided in the Supporting Text B.

### 6.3 Cryo-EM ensemble optimization software

Our method is implemented using cryoJAX [44], a JAX- and Equinox-based forward model for cryo-EM, which provides access to the JAX ecosystem, automatic GPU compilation, and efficient gradient computation [39, 56, 66]. CryoJAX offers a flexible API for defining cryo-EM image formation models, reading and manipulating RELION datasets [68], and analyzing cryo-EM datasets. We run MD simulations through OpenMM’s Python API [40], and use MDTraj [69] for communication between JAX and OpenMM.

### 6.4 Benchmark systems

#### 6.4.1 Rectangular toy model

The rectangular toy model, introduced in section 4.1, consists of four typeless atoms forming a rectangle in the *xy*-plane, with the *z*-coordinate for all atoms being zero. As mentioned earlier, this model enables us to define different force fields for the physics-based projection step with ease. Additionally, this model can be easily represented in two dimensions by its height and length, facilitating the visualization of walkers during ensemble optimization iterations. This provides valuable intuition on the ensemble optimization process. Below, we describe how we defined the force fields used for the various tests in Figure 2, as well as how the images for these tests were generated.

We define a rectangle, *R*(*h, l*) ∈ R^3×4^, through its height (*h*) and length (*l*), as a concatenation of the coordinates of four atoms placed at each corner of a rectangle, such that the rectangle’s center is at the origin.

##### Rectangle prior force field

We define the rectangle force field used for the physics-based projection in a way that only rectangular conformations are feasible. In addition, we add a prior on the feasible rectangles in the form of barriers and wells at specific conformations to mimic the behaviour of force fields for biological systems. The wells make some feasible conformations more likely, while the barriers make them less likely, but not unfeasible. We implement this in a force field defined as:

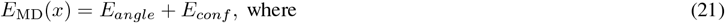

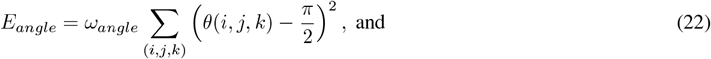

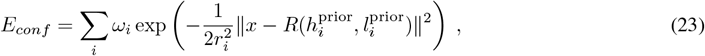

where the first term (feasibility) accounts for the angle force field for all possible successive triplets of atoms in the rectangle *R*(*h, l*), *θ*(*i, j, k*) denotes the angle between them, and *ω*_*angle*_ determines the strength of the energy term. The second term (wells and barriers prior) defines the attractive or repulsive potentials that define the prior in conformational space, where 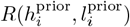 are the modes of each potential. The sign of constants *ω*_*i*_ determines if the potential is a barrier (positive) or a well (negative), and their magnitude |*ω*_*i*_| determines the strength of the prior potential (shown as dashed lines in Figure 2C-E). The constant *r*_*i*_ controls the effective radius of the wells and barriers.

For the results presented in Figure 2, we set *ω*_*angle*_ = 100 kJ mol^−1^, and consider three different combinations for the second term in the force field. In Figure 2B, we set *ω*_*i*_ = 0 kJ mol^−1^ ∀*i*, i.e., we only consider angle energy and all rectangles are equally likely. In Figures 2C-D, we define two wells centered at rectangles (32, 64) Å and (64, 64) Å with equal amplitude *ω*_1_, *ω*_2_ = −100 kJ mol^−1^, and radius 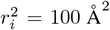; and a barrier at (48, 48) Å with *ω*_3_ = 500 kJ mol^−1^ with radius 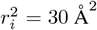. Finally, in Figure 2E, we convert the barrier at (48, 48) Å into a well with amplitude *A*_3_ = −200 kJ mol^−1^, and radius 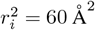.

##### Rectangle image distribution

We generate a heterogeneous set of rectangular atomic conformations to create the synthetic images used in our tests. To achieve this, we sample 800 heights and lengths from a multimodal distribution, use them to generate rectangles, and then create one image per sample as shown in Figure 2A. We define the multimodal distribution used to sample the heights and lengths as a mixture of three Gaussians with centers 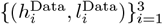 and variance 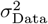, equal for all Gaussians. We determine how many images are generated from each Gaussian by assigning each a weight 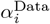. We selected the centers of the Gaussian to be (32, 64), (64, 64), and (32, 32) Å, with state populations of 0.125, 0.250, and 0.625, respectively.

##### Rectangle images

For each rectangle drawn from the GMM conformational distribution, an image is generated using the forward model defined in Eq. (6), with all rotation angles and shifts set to 0. The parameters of the projection operator (Eq. (13)) are defined as *A*_*i*1_ = 1, *A*_*ij*_ = 0 for *j*≠ 1, and *B*_*i*1_ = 128*π*^2^, *B*_*ij*_ = 1.0 for *j* 1. The parameter *B*_*i*1_ was set arbitrarily to obtain images in which the atoms could be easily visualized. The box size is set to 32, and the pixel size is set to 4 Å. The CTF is set to the identity operator, and the noise variance is defined such that the SNR is 0.5. Examples of low-pass filtered images are shown in Figure 2A.

##### Projecting onto the physical prior for the Rectangle Toy System

We perform the physics-based projection (step D in Figure 1) by running steered overdamped Langevin dynamics

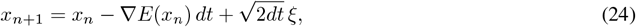

with *ξ* ∼ 𝒩 (0, 1) and

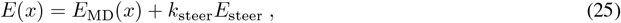

where *E*_MD_(*x*) is the force field defined in Eq. (23) and the second term defines the steering force (see Section 3.4). For these tests, we implemented an in-house version of overdamped Langevin dynamics that relies purely on JAX and Equinox. This step is analogous to the steered MD step performed for the biological benchmarks (Alanine, GroEL, and Hsp90).

##### Rectangle Ensemble Optimization

For each test (Figure 2), we run the method for 100 iterations, where each iteration cycle consists of i) two optimization steps (steps B and C) with a batch size of 20 images and a step size of 2.0 Å, and ii) 100 steps of steered overdamped Langevin dynamics with *dt* = 0.001, and *k* ^steer^ = 1000.0 kJ mol^−1^ Å^−2^ (step D). For this simplified system, no alignment is necessary since we simulate the images without rotation.

#### 6.4.2 Alanine Tri-peptide

For the next benchmark, we use the tri-alanine benchmark system (section 4.2). This is a 42-atom system that we use to test our pipeline when the projection step (Figure 1D) is performed via steered MD. With this system, we demonstrate the importance of the physical prior, not only for the structural stability of the walkers but also for robustness against overfitting to image artifacts. We generate the images by sampling conformations 3Ala-A and 3Ala-B with probabilities 0.7 and 0.3, respectively, and applying the forward model defined in Eq. (6) with an SNR of 0.5. We generate a total of 5,000 images across the two conformations. In all cases, the optimization step is performed for the non-hydrogen atoms. We initialize the algorithm with both walkers starting from a third conformation, 3Ala-C, and use 3Ala-A for alignment (see section 6.2.1). We conducted two tests: in the first, we optimized without a physical prior by skipping the projection step in the pipeline (Step D in Figure 1); in the second, we optimized with a physical prior by running all the steps. In both cases, we run our method for 100 complete iterations. For the case with a prior, each iteration cycle consists of i) alignment to 3Ala-A (step A), ii) ten optimization steps (steps B and C) with a batch size of 500 images and a step size of 0.25 Å, and iii) 500 steps of steered MD (step D). For the case without a prior, we i) align to 3Ala-A (step A), and ii) perform 20 optimization steps (steps B and C) with a batch size of 500 images and a step size of 0.1 Å. MD simulations are performed with OpenMM [40] with a time step of 2 *fs*. The force constant was set to *k* = 3 × 10^4^ kJ mol^−1^ Å^−2^. The MD simulations are run in vacuum, using the Amber 14 force field [70]. The optimized structures are shown in Figure 3C for the case with a physical prior (blue, left) and without one (orange, right); the reference structures are shown in gray. Because the cryo-EM optimization step is performed for the non-hydrogen atoms, we only show these atoms in Figure 3C.

#### 6.4.3 Chain A of GroEL

We benchmark the method using an ensemble constructed from chain A of the chaperonin GroEL in two conformations, as described in [23]. The first structure is obtained by extracting chain A from the apo-GroEL structure (PDB ID: 1XCK). The second structure is obtained by constructing a homology model from chain A of GroEL-ADP in the relaxed allosteric state (PDB ID: 4KI8) using the apo-GroEL sequence. We refer to these structures as apo-GroEL and holo-GroEL, respectively. Each atomic model comprises 7,835 atoms. We use this system to assess method performance under various sources of uncertainty in the cryo-EM dataset.

We benchmark the method against increasing noise in the cryo-EM image particles. To do this, we simulate 1000 images from holo-GroEL at different SNR values (10, 1.0, 0.1, and 0.01). For each SNR value, we run our method for 100 iterations, each consisting of i) 10 optimization steps (steps B and C) with batch size of 500 images and a step size of 2.0 Å, and ii) 500 steered MD steps (Step D) with a step size of 2 *fs*, in vacuum and force constant *k* = 3 × 10^5^ kJ mol^−1^ Å^−2^. We compare this to an “ideal case”, where we run a steered MD simulation using the same force constant and the true holo-GroEL structure as a reference for the biasing force. For this case, we ran the steered MD simulation for 500, 000 steps to match the aggregated MD time run during the ensemble optimization.

In the second set of tests, we benchmark our method against inaccuracies in the pose parameters. Heterogeneity methods typically assume that the pose information is known, often derived from a consensus reconstruction. In this test, we examine how this assumption influences the performance of our method. We generate 20, 000 images from apo-GroEL and holo-GroEL with probabilities of 0.7 and 0.3, respectively; defocus values between 1.0 *µm* and 1.5 *µm*, and SNR = 0.1. We estimate the poses by running *ab initio* reconstruction and homogeneous refinement on this dataset. We then align the apo-GroEL model with the resulting consensus volume and use it to initialize the structural ensemble. We run our method for 100 iterations, each consisting of i) global RMSD alignment to apo-GroEL and local rigid-body alignment to the consensus volume (step A), ii) 10 optimization steps (steps B and C) with a batch size of 500 images, and a step size of 2.0 Å, and iii) 500 steered MD steps with a force constant of *k* = 3 × 10^5^ kJ mol^−1^ Å^−2^ (step D). We repeat the test using the known poses, keeping all other parameters identical for comparison.

#### 6.4.4 Real data of Hsp90

We benchmark the method by applying it to real data of the hsp90-p23-GR dataset [54] (EMPIAR-11028). This dataset challenges our method with real experimental data, initialization from a distant atomic structure, and model misspecification. We focus on the hsp90-p23 subset assigned in [54]. The associated deposited structure (PDB ID: 7krj) shows hsp90 with its two arm-like subunits (chains A and B) in a closed state, forming a compact V shape. In [55], a heterogeneous cryo-EM dataset for hsp90 was designed by generating two motions, one producing a structure where the chains open to form a wide V-shape (Figure 5A). We select as the initial structure the model with the largest deformation along this motion, which has a 10 Å heavy-atom RMSD from the hsp90 macromolecule in the deposited PDB from [54] (PDB 7krj), estimated from the consensus reconstruction volume (EMD-23006, 2.6 Å in resolution). Beyond this poor initialization, the structure also suffers from model misspecification due to the absence of the p23 protein. We prepare the initial model for simulation using PDBFixer [71]. Hereafter, we refer to this initial structure as open-hsp90.

The hsp90-p23-GR dataset includes micrographs, the locations of the picked particles within each micrograph, and estimated pose and CTF parameters. We extract the particles using the locations provided by the dataset. We downsampled to a box size of 128 pixels for computational efficiency and estimated the poses through consensus reconstruction with CryoSPARC [53], resulting in a total of 145, 069 particles. Typically, during the consensus reconstruction process, the dataset is split in half; we designate half-1 as the training dataset and half-2 as the test dataset. We use the half-volume associated with the training dataset for the local alignment step described in section 6.2.1.

We initialize our method with one walker assigned 100 % of the weight from open-hsp90 and run it for 100 iterations. Each iteration consists of i) global RMSD alignment to open-hsp90 and local rigib-body alignment to the half-volume associated to the training images (step A), ii) 10 optimization steps (steps B and C) with a batch size of 250 images, and a step size of 2.0Å, and iii) 1000 steered MD steps with a force constant of *k* = 3 × 10^5^kJ mol^−1^Å^−2^ and a step size of 2 *fs* (step D). The total runtime is 18 minutes on an NVIDIA A100 GPU with 40 GB of memory. The induced conformational change corresponds to an optimized structure with a 10Å heavy-atom RMSD to the initial structure.

## Data availability

The cryo-EM ensemble optimization method repository can be accessed at https://github.com/flatironinstitute/cryojax-ensemble-optimization, and the necessary data – including atomic models, config files, and instructions – to reproduce the results from the different benchmark systems are available in https://zenodo.org/records/17420048.

## Acknowledgments and Disclosure of Funding

The authors thank Michael O’Brien, Miro Astore, Lars Dingeldein, Wai Shing Tang, Aaditya Rangan, Geoff Woollard, Daniel Needleman, Sonya Hanson, and Niko Grigorieff for helpful discussions. The Flatiron Institute is a division of the Simons Foundation. The work was supported by NIH/NIGMS (R01GM136780 and 1R35GM157226), the Alfred P. Sloan Foundation (FG-2023-20853), AFOSR (FA9550-21-1-0317), and the Simons Foundation (1288155).

## A Supporting Information

### A.1 Image-to-structure likelihood computation

In this section, we provide additional details for the quantities required to compute the image-to-structure likelihood.

#### A.1.1 Scale and offset

As described in section 6.1.3, the optimal scale and offset for each image are obtained by directly optimizing

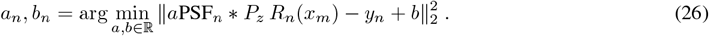

The analytical solution can be obtained by direct optimization and is given by

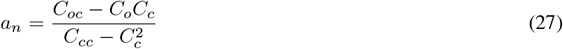

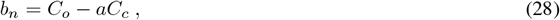

where the quantities *C*_*o*_, *C*_*c*_, *C*_*oc*_, and *C*_*cc*_ are defined as:

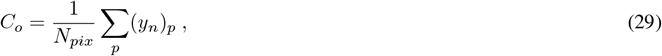

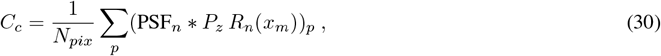

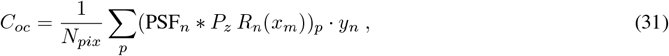

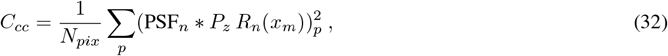

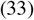

where the sum is over all pixels *p* = 1, …, *N*_*pix*_.

#### A.1.2 Marginalization of the noise variance

In section 6.1.3, we simplify the image-to-structure likelihood by marginalizing over the noise variance. We assume that the noise is Gaussian white noise, which can be achieved by whitening the cryo-EM images [72]. The marginalization is achieved by integrating over the noise variance, assuming it follows a probability distribution *p*(*σ*^2^):

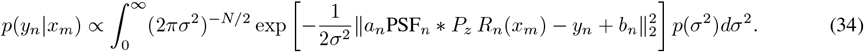

We model the distribution *p*(*σ*^2^) using Jeffrey’s prior [63, 64] for the variance of an isotropic multivariate Gaussian distribution, which we define below.

#### Jeffrey’s prior for the variance of a multivariate isotropic Gaussian distribution

Jeffrey’s prior is typically used in Bayesian statistics to define a non-informative prior on a parameter or, more generally, a parameter space. For a parameter *θ*, this prior is defined as the square root of the determinant of the Fisher information:

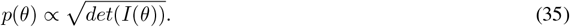

For an N-dimensional multivariate Gaussian distribution with isotropic variance, *N* (*µ, σ*^2^*I*_*N*_ ), the Fisher information can be computed as:

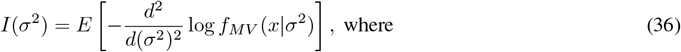

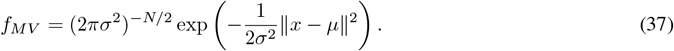

First we take the second derivative of log *f*(*x*|σ^2^ ) with respect to σ^2^ 804 yielding:

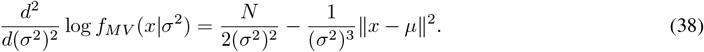

The computation of the expected value can be performed using the linearity of expectation, and the fact that the covariance matrix is isotropic, and thus each entry of the random variable *x* ∼ *N* (*µ, σ*^2^*I*_*N*_ ) is independent and identically distributed: *x*_*i*_ ∼ *N* (*µ*_*i*_|*σ*^2^). Using these two facts, we obtain that the Fisher information is:

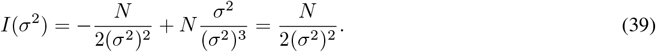

Finally, we obtain that Jeffrey’s prior for *σ*^2^ is

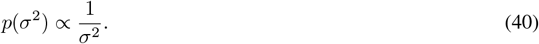

#### Using Jeffrey’s prior to marginalize the image-to-structure likelihood

Using the prior for the variance derived in Eq. (4 ), the marginal image-to-structure likelihood is

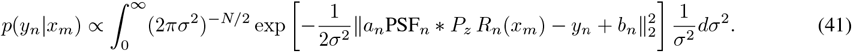

This expression can be related to the following integral:

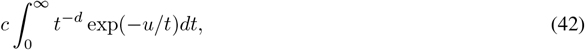

by the defining the constants as 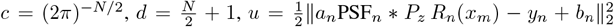, and the integration variable as *t* = *σ*^2^. This integral can be solved by changing variables as 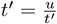, which gives us a Gamma integral with solution:

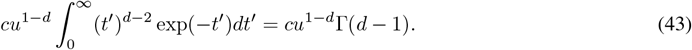

Therefore, the marginalized likelihood is

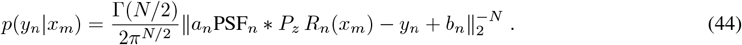

## B Scaling factor of the biasing force in the prior projection step

In this section, we show how to compute the scaling factor used in the biasing force for the projection step, as defined in section 6.2.4. We define the scaling factor, *k*_steer_, as the solution to the following equation

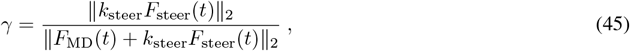

where *γ* is the desired proportion of the contribution of the steering force to the total force at frame *t*, and ∥ · ∥ is the *ℓ*2-norm. This parameter is defined by the user, and is achieved by modifying *k*_steer_. That is, we solve for *k*_steer_ given a value for *γ*.

To solve for *k*_steer_, first take the square of the previous equation to obtain:

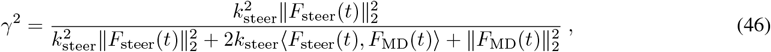

where ⟨·,·⟩ denotes the Euclidean inner product. The value of *k*_steer_ is obtained by solving the following quadratic equation at each frame *t*:

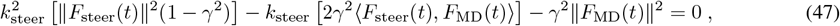

With

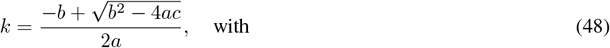

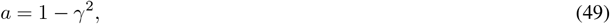

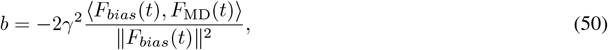

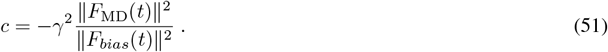

## C Structure-to-volume forward model

We define the structure-to-volume forward model (*F*_*V*_ ) by extending the projection operator defined in section 6.1. The generated volume is discretized in a 3D grid with entries:

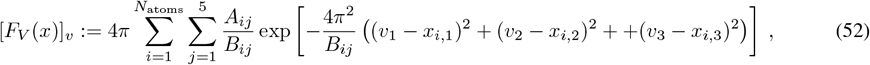

\where *v* = (*v*_1_, *v*_2_, *v*_3_) corresponds to the voxel coordinates in the 3D grid. The quantities *A*_*ij*_ and *B*_*ij*_ are the electron form factors for each atom (see section 6.1), and 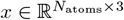 are the atomic coordinates.

## D Geometry analysis of obtained structures

### D.1 Rectangular toy model

We measure the quality of the rectangular structures obtained in section 4.1 in two different ways. For the first metric, for all walkers, we compute the angles at each corner of the rectangle 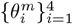, and compute the root mean square error (RMSE) with respect to a 90^◦^ angle, that is

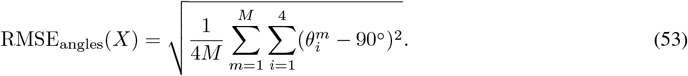

The second metric involves computing the average ratio between the diagonals of the rectangle defined by the walkers, which should be 1.0 for a rectangular structure. Table 1

**Table 1.** Geometry metrics for the rectangles obtained in section 4.1. **Left**: Walkers used to compute the metrics identified by the panels in Figure 2. **Middle**: Mean squared error between the angles computed at each corner and an angle of 90^◦^ averaged over all walkers. **Right**: Ratio between the diagonals of the obtained structures. A value of 1.0 means that the structure is rectangular.

| Walkers | RMSE<sub>angles</sub> | Ratio between diagonals |
| --- | --- | --- |
| Figure 2B | 4.0° | 0.98 |
| Figure 2C | 3.64° | 0.98 |
| Figure 2D | 3.15° | 1.0 |
| Figure 2E | 3.16° | 1.01 |

The metrics obtained for the sets of walkers shown in Figure 2 are summarized in Table 1. These results demonstrate that the physical prior designed for this system results in structures that closely match a rectangular geometry.

## Notes

### Competing Interest Statement

The authors have declared no competing interest.

